# Environment driven conformational changes modulate H-NS DNA bridging activity

**DOI:** 10.1101/097436

**Authors:** Ramon A. van der Valk, Jocelyne Vreede, Geri F. Moolenaar, Andreas Hofmann, Nora Goosen, Remus T. Dame

## Abstract

Bacteria frequently need to adapt to altered environmental conditions. Adaptation requires changes in gene expression, often mediated by global regulators of transcription. The nucleoid-associated protein H-NS is a key global regulator in *Gram*-negative bacteria, and is believed to be a crucial player in bacterial chromatin organization via its DNA bridging activity. H-NS activity *in vivo* is modulated by physico-chemical factors (osmolarity, pH, temperature) and interaction partners. Mechanistically it is unclear how functional modulation of H-NS by such factors is achieved. Here, we show that a diverse spectrum of H-NS modulators alter the ability of H-NS to bridge DNA. Changes in monovalent and divalent ion concentrations drive an abrupt switch between a bridging and non-bridging DNA binding mode. Similarly, synergistic and antagonistic co-regulators modulate the DNA bridging efficiency. Structural studies suggest a conserved mechanism: H-NS switches between a “closed” and an “open”, bridging competent, conformation driven by environmental cues and interaction partners.

---

The bacterial genome is organized and compacted by nucleoid-associated proteins^1–7^. A key protein in nucleoid organization of Gram-negative bacteria is the Histone-like Nucleoid Structuring protein (H-NS). Genome-wide binding studies have revealed that H-NS binds along the genome in long patches^8–12^, which have been proposed to mediate the formation of genomic loops^13,14^. H-NS is also an important regulator of global gene expression, implied in mediating global transcriptional responses to environmental stimuli (osmolarity, pH, temperature)^15^. A large fraction of *Escherichia coli* and *Salmonella* genes (5-10%) is regulated (usually repressed) by the action of H-NS. H-NS operation is modulated by environmental stimuli and through interplay with other proteins ^15,16^. In solution the H-NS protein exists as a dimer, which oligomerizes at high concentrations^17,18^. H-NS consists of three structural domains: a C-terminal domain responsible for DNA binding^19^, a N-terminal dimerization domain^20–23^ and a central dimer-dimer interaction domain responsible for multimer formation^24,25^. These two interaction domains are connected by a long α- helix^24^. H-NS exhibits two seemingly distinct DNA binding modes: DNA bridging^26–30^, the condensation of DNA by intra- and inter- molecular DNA binding by H-NS and DNA stiffening, the rigidification of DNA through the formation of a rigid H-NS-DNA filament ^31–33^. These modes have been attributed to the basic functional H-NS unit (a dimer) binding to DNA either *in cis* or *in trans*^34,35^. H-NS paralogues StpA, Sfh, Hfp, and truncated derivatives such as H-NST, have been proposed to modulate H-NS function by forming heteromers with H-NS^36–40^, with DNA binding properties different from homomeric H-NS. Members of the Hha/YmoA family of proteins are H-NS co-regulators with limited sequence homology to H-NS ^41^. At many targets along the genome, H-NS and Hha co-localize. Localization of Hha at these sites is strictly H-NS dependent, whereas the genome-wide binding pattern of H-NS is only mildly affected by Hha^42^.

Although evidence has been put forward that the concentration of divalent ions determines the binding mode of H-NS^33^, a mechanistic explanation is lacking. Moreover, the possible effect of co-regulators of H-NS, such as Hha, on these binding modes has remained unexplored. To obtain a better understanding of the molecular basis underlying the H-NS binding modes, it is crucial to determine the effects of ion valence and concentration, as well as the presence of helper proteins on both the stiffening and the bridging mode. Here we investigate DNA stiffening on short DNA tethers using Tethered Particle Motion (TPM). As *intramolecular* DNA bridging does not occur on short DNA tethers, DNA stiffening can be uncoupled from *DNA bridging*. In addition, to accurately determine *intermolecular* DNA bridging efficiencies in solution, we developed a sensitive quantitative bulk assay. Using these two assays we unravel the assembly pathway of bridged DNA-H-NS-DNA complexes and the roles of mono- and divalent ions, helper proteins Hha and YdgT, and truncated H-NS derivatives. Finally, Molecular Dynamics (MD) simulations reveal that ions and interacting proteins *directly* alter H-NS structure from a “closed” bridging incapable to an “open” bridging capable conformation, thus providing a molecular understanding of the modulation of H-NS function.

## RESULTS

### The role of Mg^2+^ and H-NS multimerization in DNA bridging and DNA stiffening

In order to dissect the role of divalent ions in the formation of bridged and stiffened complexes, we applied a novel, sensitive *quantitative* DNA bridging assay and carried out TPM experiments (providing a quantitative and selective readout of DNA stiffening). The DNA bridging assay relies on immobilization of bait DNA on magnetic microparticles and the capture and detection of ^32^P labeled prey DNA if DNA-DNA bridge formation occurs. 80% of initial prey DNA is recovered at high H-NS concentrations (see figure 1a). In the absence of either H-NS protein or bait DNA, no prey DNA is recovered under our experimental conditions. Next, we used this assay to quantify the DNA bridging efficiency of H-NS as a function of the amount of Mg^2+^ ions (see figure 1b), reproducing the qualitative results of Liu *et al*^33^ and providing independent confirmation of the previously observed effects. The concentration range from 0 – 10 mM Mg^2+^ is considered to be physiologically relevant.^43^. Importantly, the transition from no bridging to complete bridging is abrupt between 4 – 6 mM Mg^2+^, indicating that changes in Mg^2+^ concentration might drive a binary switch.

**Figure 1.**
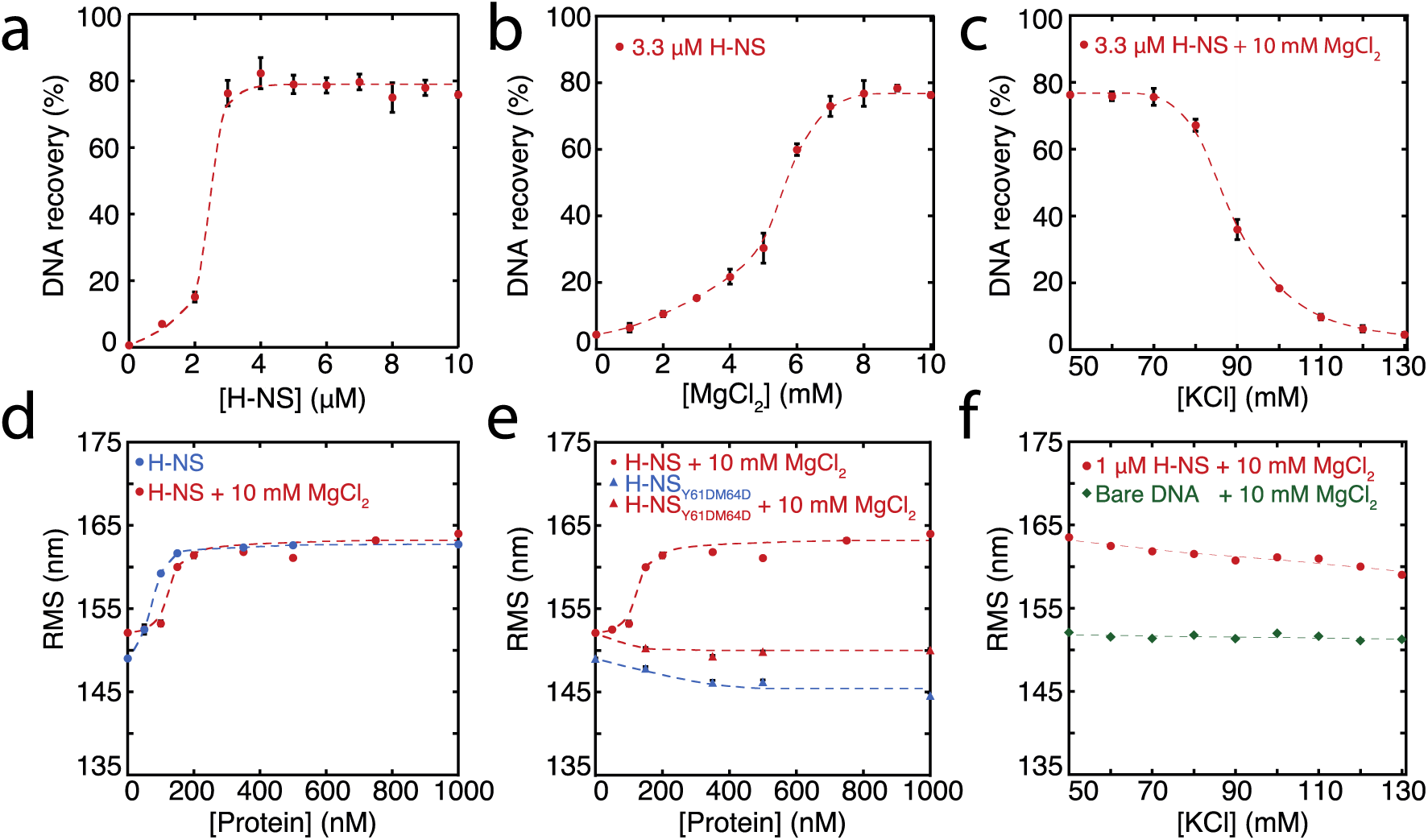
Modulation of H-NS function by ionic conditions. (a) DNA bridging efficiency as a function of H-NS concentration in the presence of 10 mM MgCl_2_. (b) DNA bridging efficiency as a function of MgCl_2_ concentration. (c) DNA bridging efficiency as a function of KCl concentration. (d) Root Mean Square displacement (RMS) as a function of H-NS concentration, in the presence and absence of 10 mM MgCl_2_ (N>70, per data point). (e) RMS of DNA as a function of H-NS_Y61DM64D_ in the presence and absence of MgCl_2_ (N>70, for each point). (f) Extension of DNA as a function of KCl concentration (N>70, for each point). Error bars indicate standard deviation. Dashed lines are to guide the eye.

Earlier studies have shown that H-NS binding along a single DNA molecule results in DNA stiffening^31,33^. However, due to the co-occurrence of DNA bridging, previous studies were incapable of measuring DNA stiffening in the presence of Mg^2+^. Here we used TPM to investigate DNA stiffening as a function of Mg^2+^ concentration. In TPM, the Root Mean Square displacement (RMS) of bead movement is a direct reflection of tether stiffness and length. DNA binding proteins can affect both, but previous studies have shown that the DNA contour length is not affected by binding of H-NS^26,28^. Thus an increase in stiffness due to H-NS binding translates into a higher RMS value of a DNA tether (figure 1d). Here we measured the effects of H-NS on DNA stiffness in the absence and presence of 10 mM Mg^2+^ and confirmed that H-NS stiffens DNA^31,33^; importantly our experiments reveal that Mg^2+^ does not affect the stiffness of the fully formed H-NS-DNA complexes at saturation, as in both conditions the RMS is the same (figure 1d). Analysis of the binding characteristics using the McGhee-von Hippel equation revealed that the association constant of H-NS is somewhat reduced in the presence of Mg^2+^, while cooperativity increases under these conditions (Figure 3 - figure supplement 1a,d). The reduction in DNA binding affinity of H-NS may be attributed to shielding of the negatively charged phosphate backbone by Mg^2+^.

The cooperative binding of HNS and DNA stiffening observed by TPM suggest that H-NS multimerizes along DNA, likely via the recently defined dimer-dimer interaction domain^24^. Multimerization along DNA has been previously suggested^22,39^, but has never been conclusively demonstrated. To test this hypothesis, we generated a mutant, H-NS_Y61DM64D_, predicted to have disrupted dimer-dimer interaction based on the H-NS_1-83_ crystal structure^24^. Size exclusion chromatography showed that this H-NS mutant indeed exists solely as a dimer in solution independent of protein concentration (Figure 1 - figure supplement 1b), whereas wild type H-NS forms large multimeric structures (Figure 1 - figure supplement 1a and ^24,25^). The multimerization behavior of both proteins was unaffected by the presence of Mg^2+^. Electrophoretic Mobility Shift Assay confirmed that the DNA binding of the H-NS mutant is intact (figure 1 - figure supplement 3). TPM experiments reveal that H-NS_Y61DM64D_ binding does not lead to the formation of stiff H-NS-DNA filaments (figure 1e). The RMS is reduced compared to that of bare DNA, indicating not only that dimer-dimer interactions are disrupted, but also that individual H-NS dimers mildly distort DNA. DNA bridging experiments reveal that H-NS_Y61DM64D_ is also incapable of forming DNA-H-NS-DNA complexes (Figure 1 - figure supplement 2). This indicates that individual H-NS_Y61DM64D_ dimers do not form stable bridges and that dimer-mediated bridging, involving H-NS-dimer binding cooperativity due to high local DNA concentration adjacent to existing bridges^26,27^, is insufficient to explain the formation of bridged DNA-H-NS-DNA complexes. The nature of the effect of Mg^2+^ on DNA bridging efficiency is not understood. If the positive effect of Mg^2+^ would be solely due to it facilitating interactions between the DNA phosphate backbone and negatively charged residues on H-NS, an increase in affinity would be expected, but this is not observed (figure 3 - figure supplement 1). As Mg^2+^ does not affect the multimeric state of H-NS in solution (Figure 1 - figure supplement 1), a model involving an effect on H-NS multimerization can be excluded. A structural effect of Mg^2+^ on individual units within H-NS filaments could explain the observed effects of Mg^2+^ on the bridging efficiency of H-NS.

### Mg^2+^ alters H-NS structure

To investigate the role of Mg^2+^ on individual H-NS dimers we carried out MD simulations of an H-NS dimer in both the absence and presence of Mg^2+^, using our previously established model of a full-length H-NS dimer^14^. Visual inspection of the H-NS dimer simulations at 50 mM KCl reveals that H-NS changes from an “open” extended conformation into more compact “closed” shapes (see figure 2a for snapshots from these simulations or figure 2 - figure supplement 2 for a movie of one such simulation). The three domains in H-NS interact, and these inter-domain interactions are facilitated by partial unfolding and buckling of the long central α helix (helix α3) connecting the dimerization and dimer-dimer interaction domains. By computing the average distance between the donor and acceptor of the hydrogen bonds within helix α3, denoted as <d_O-H_>, we determined that the buckle forms in region Glu42-Ala49 (figure 2b), as <d_O-H_> exceeds the hydrogen bond distance threshold of 0.35 nm. By analyzing the O-H distance between residues Ser45 and Ala49 in time, key residues at the site of buckle formation (see Figure 2 - figure supplement 3), we found that buckles can be reversible and irreversible, within the simulation time scale of 50 ns. Reversible buckles, caused by thermal fluctuations, typically last a few nanoseconds and occur several times during a single simulation run (see green line in Figure 2 - figure supplement 3 for an example). Irreversible buckles, stabilized by inter-domain interactions, do not return to a helical conformation during our simulations (see red line in Figure 2 - figure supplement 3 for an example). To characterize the interactions that occur during the simulations, we generated contact maps that show the probability of finding interactions between residues. A contact is counted if the minimum distance, calculated every 10 ps, between a residue pair is 0.6 nm or less. To get the probability, these counts are normalized over the total simulation time (Figure 2 - figure supplement 1a). These maps reveal that the DNA binding domain interacts with other parts of the protein complex. In absence of Mg^2+^, the DNA binding domain interacts with the dimerization domain, rendering the DNA binding QGR motif (residues 112-114) ^44^ of one DNA binding domain inaccessible (see the snapshots in figure 2a). Even though the simulations were performed in absence of DNA, these observations indicate that DNA bridging is no longer possible in such a conformation, as the H-NS dimer can bind DNA only through its remaining/second DNA binding domain. In this ‘closed’ conformation, interactions occur between the IRT residues at position 10-12 and the AMDEQGK residues at position 122-128. These interactions are hydrophilic in nature, supplemented by a salt bridge between R11 and D124 or E125. In the presence of Mg^2+^ interactions between the DNA binding domain and the dimerization domain no longer occur (see the snapshots in figure 2a or the movie in Figure 2 - figure supplement 4), The absence of such interactions is further illustrated by the contact map in Figure 2 - figure supplement 1b and in higher detail in Figure 2 - figure supplement 6. The likelihood of finding Mg^2+^ interacting with (i.e. being within 0.6 nm of) H-NS residues, indicated by P_Mg2+_, revealed that Mg^2+^ has a preference for glutamate residues in region 22-35 (figure 2c), where the ions shield this region from interacting with the DNA binding domains. The magnesium ions transiently interact with the glutamate residues, with residence times in the order of a few ns (as illustrated in Figure 2 - figure supplement 4) The presence of Mg^2+^ stabilizes the “open” conformation of H-NS, ensuring that DNA bridging can occur. Furthermore, we noted that Mg^2+^ is recruited to the location of the buckle and may directly stabilize helix α3 through interactions with Glu42 (Figure 2 - figure supplement 8), resulting in an “open”, bridging capable, H-NS conformation (see Figure 2 - figure supplement 2 and 4). These data suggest that Mg^2+^ modulates H-NS by shielding interactions between the DNA binding domain and dimerization domain, and by influencing the conformation of helix α3.

**Figure 2.**
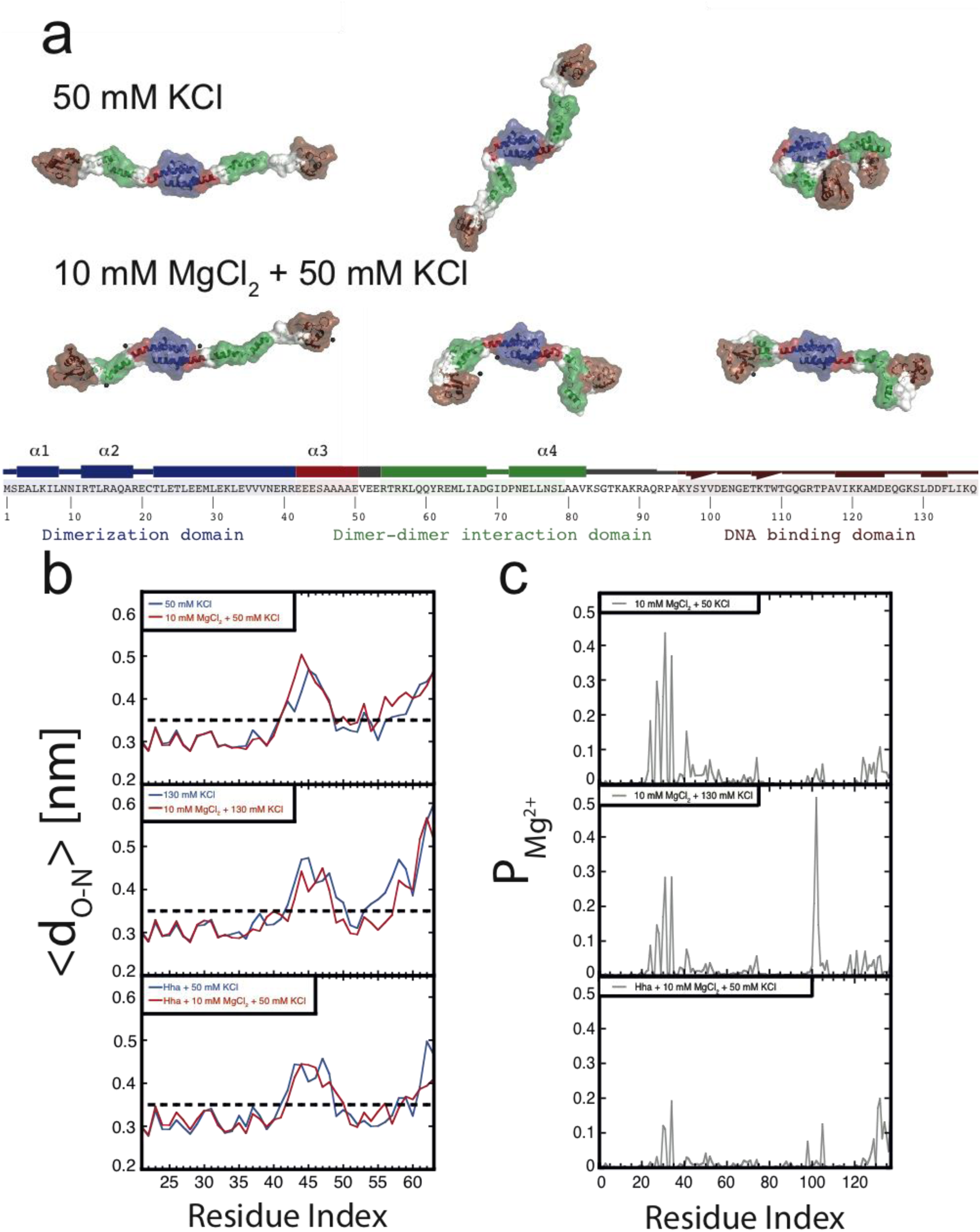
Conformation of the H-NS dimer as a function of osmolarity. (a) Snapshots depicting representative conformation of H-NS in the simulations with 50 mM KCl (top) and 10 mM MgCl_2_ + 50 mM KCl (bottom) (For the full movies depicting these effects see Figure 2 - figure supplement 2 and 4). The buckle region is highlighted in red and the domains are colored blue, green and brown for the dimerization domain, the dimer-dimer interface and the DNA binding domain respectively. The conserved motif involved in DNA binding is highlighted as ball-stick models. Mg^2+^ ions are shown as dark gray orbs. The protein is shown in ribbon representation with a transparent surface. The atomic radii in the protein were set to 3 A to smooth the surface. (b) Location of buckle in helix α3. The average distance d_O-N_ between donor and acceptor in the helical hydrogen bond in helix α3 is plotted as a function of the residue index of the acceptor. The dashed black line in the graphs indicates the distance threshold for forming a hydrogen bond. Time traces of these distances are given in Figure 2 - figure supplement 3. (c) Location of Mg^2+^ on H-NS. The probability of finding Mg^2+^ ions within 0.6 nm of an H-NS residue, P_Mg^2+^_, is plotted as function of the residue index for the three systems containing Mg^2+^.

### Modulation of DNA bridging by osmotic factors

Although it has long been known that the expression of some H-NS controlled genes (such as the *proU* operon) is modulated by the osmolarity of the medium^45^, the underlying mechanism remains undetermined. Previous studies have revealed that the H-NS DNA stiffening mode is mildly sensitive to the KCl concentration^31,33^. Using TPM we were able to confirm these observations. The reduction in DNA stiffening is gradual^34^ and modest (figure 1f). It is thus questionable whether the multimerization of H-NS along DNA alone is sufficient to explain its role in repression of transcription (and modulation thereof). Could the modulation of gene repression be due to ionic effects on DNA bridging efficiency? Using our DNA bridging assay we observed complete abolishment of H-NS DNA bridging by KCl at concentrations exceeding 120 mM (figure 1c), a binary response, similar to what we observed for the Mg^2+^ titration. This *in vitro* observation mirrors the *in vivo* response of the ProU operon, at which KCl concentrations exceeding 100 mM are required to alleviate H-NS repression ^45^. This strong and abrupt effect on DNA bridging, while leaving DNA stiffening essentially unaffected, might indicate that H-NS reverts to the “closed” conformation by the addition of KCl. To investigate this effect at a structural level we performed MD simulations at high KCl concentrations. The presence of 130 mM KCl alters interactions between the various domains in H-NS (see contact maps in Figure 2 - figure supplement 1c and d). Figure 2 - figure supplement 5 shows the likelihood of finding interactions between K^+^ and Cl^-^ ions and H-NS residues, denoted as P_k+_ and P_cl-_ at high and low KCl concentrations, and in absence and presence of Mg^2+^. K+ ions have a higher probability to interact with (negatively charged) residues in the dimerization domain and the DNA binding domain (Figure 2 - figure supplement 5). The pattern of interaction between H-NS and Cl^-^ ions is different at 50 and 130 mM KCl. This is most pronounced in the DNA binding domain. The average distance between the donors and acceptors in the hydrogen bonds within helix α3 (figure 2b) indicates that that buckles in helix α3 also occur at a high KCl concentration, at the same location as determined by low salt conditions. The presence of Mg^2+^ does not significantly alter the occurrence of buckles (figure 2b). Instead, Mg^2+^ is capable of deterring interactions between the DNA binding domain and the dimerization domain through recruitment to the dimerization domain (figure 2c). The probability of interactions occurring between the DNA binding domain and the dimerization domain is reduced significantly in the presence of Mg^2+^ (figure 2- figure supplement 6). This inhibits H-NS bridging by promoting the “closed” conformation of H-NS even in the presence of Mg^2+^ and elucidates the modulatory and regulatory effects of KCl and osmolarity.

### Modulation of DNA bridging and DNA stiffening by truncated H-NS variants

In addition to environmental factors such as osmolarity, H-NS activity is affected by interactions with other proteins *in vivo*. Members of the Hha/YmoA protein family, such as Hha and YdgT, are known to cooperate with H-NS in repression of genes^38^, while other proteins such as H-NST are capable of inhibiting H-NS function^35^, likely by hampering H-NS multimerization. To systematically investigate the latter mechanism, we designed and synthesized truncated H-NS derivatives, aiming to target the H-NS dimerization domain (H-NS_1-58_) or dimer-dimer interaction domain (H-NS_56-83_). Interfering with H-NS dimerization, through the addition of H-NS1-58 in DNA bridging experiments, we observed a reduction in DNA recovery from 75% to 20% at ratios higher than1: 3 H-NS_1-58_/ H-NS (figure 3a). Similarly, targeting the dimer-dimer interaction domain (through H-NS_56-82_) resulted in complete abolishment of DNA bridging (figure 3a). Next, we investigated the effects of H-NS derivatives on H-NS DNA stiffening using TPM. Only at very high inhibitor concentrations (30-fold excess of inhibitor) reduction of DNA stiffening was observed (figure 3b). These experiments reveal that both H-NS dimerization and dimer-dimer interactions can be effectively targeted for inhibition of H-NS activity and that the respective domains are crucial to the formation of bridged filaments. This strongly suggests that natural H-NS inhibitors such as H-NST operate by disrupting DNA bridging and provides clues for rational design of artificial peptide inhibitors of H-NS.

**Figure 3.**
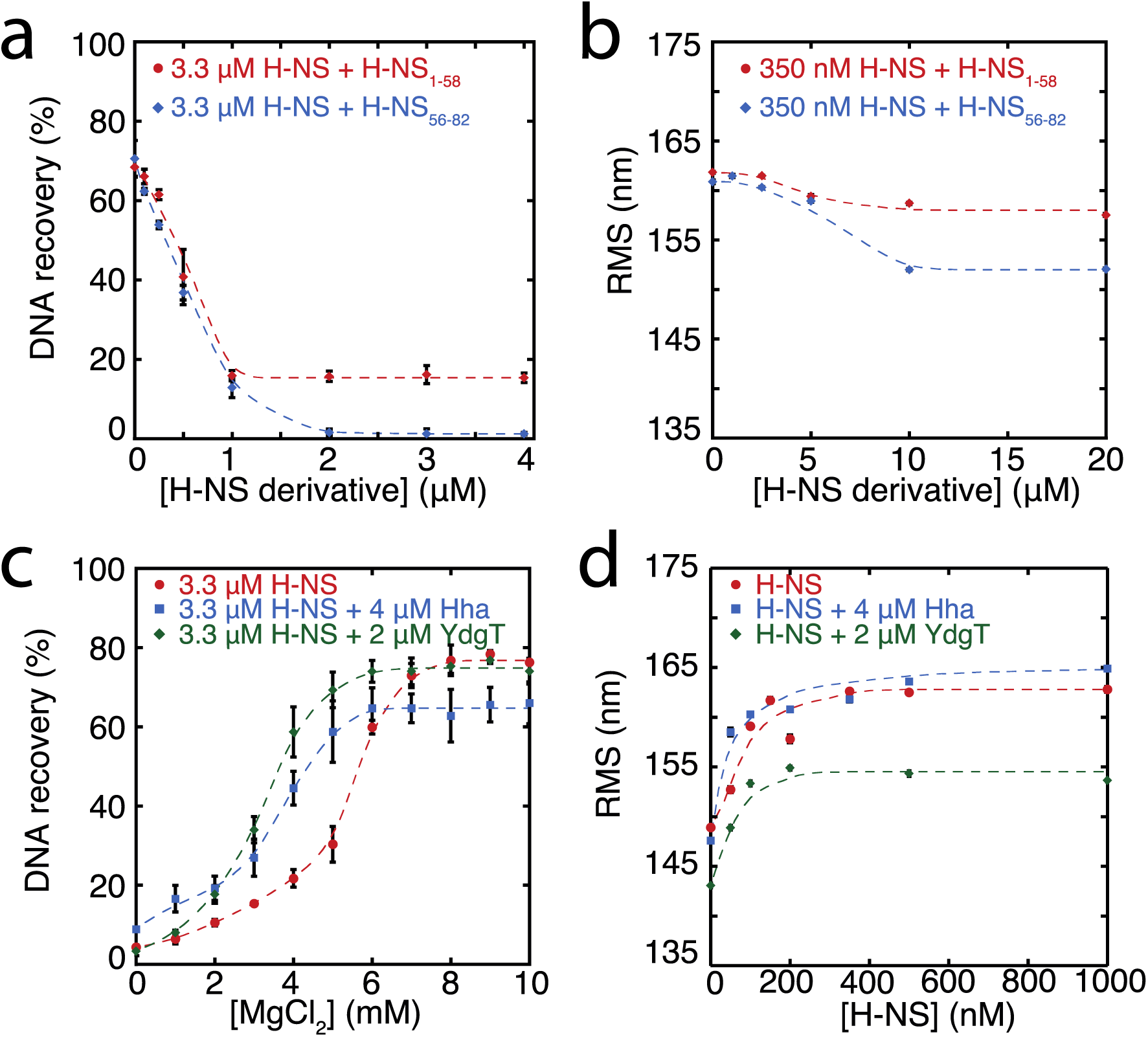
Modulation of H-NS function by protein cofactors. (a) DNA bridging efficiency as a function of inhibiting peptides targeting either the dimerization domain (H-NS_1-58_) and multimerization domain (H-NS_56-82_). (b) Root Mean Square displacement (RMS) as a function of inhibiting peptides targeting either the dimerization (H-NS_1-58_) and multimerization (H-NS_56-82_) (N>70, for each point). (c) DNA bridging efficiency as a function of Mg^2+^ concentration in the presence and absence of 4 μM Hha or 2 μM YdgT. (d) RMS of DNA in the presence of H-NS, H-NS-Hha, and H-NS-YdgT. Dashed lines are to guide the eye (N>60, for each point).

### Modulation of H-NS by Hha and YdgT

Gene regulation by H-NS often occurs in conjunction with other proteins; these co-regulators are known to interact with H-NS at specific loci along the genome. Two such proteins, Hha, and YdgT, are members of the Hha/YmoA^41^ family of proteins. In order to understand the modulation of H-NS function by these proteins we investigated their influence on the H-NS DNA binding modes. We observed that Hha, when added at equimolar concentrations, enhances DNA bridging by H-NS at low Mg^2+^ concentrations (figure 3c). A similar enhancement of H-NS mediated DNA bridging was observed with the Hha paralogue, YdgT. While Hha and YdgT promote DNA bridging to a similar extent, YdgT promotes DNA bridging at significantly lower concentrations, likely due to a higher affinity for H-NS. To determine whether enhanced DNA bridging is due to structural changes in H-NS-DNA filaments, we investigated the effects of Hha and YdgT on DNA stiffening (figure 3d). TPM experiments show a mild increase in H-NS mediated DNA stiffening in the presence of Hha. We observe a negative offset in RMS in the presence of YdgT, indicating a more compact conformation. This reduction in RMS promotes intramolecular DNA contacts; specifically this is evident in TPM experiments containing H-NS, YdgT, and Mg^2+^, where in the presence of Mg^2+^ and YdgT, H-NS causes “DNA collapse” in TPM (data not shown), similar to earlier observations for H-NS in the presence of Mg^2+^ without ^33,46^ or with Hha added ^46^. One possible explanation for the effects of Hha and YdgT, is that they effectively increase the DNA binding affinity of H-NS^47^. We find that the affinity of H-NS is significantly enhanced by Hha, but it is not significantly altered by YdgT (figure 3 - figure supplement 1d). Notably, YdgT lacks several positive residues (such as K32 and R81) present in Hha, which have been proposed to interact with DNA and to enhance DNA binding of H-NS ^47^. We also find that Hha significantly reduces the cooperativity of H-NS DNA stiffening in both the presence and absence of Mg^2+^, whereas no such effect is observed for YdgT (figure 3 - figure supplement 1). The cooperativity in binding of H-NS is attributed to filament formation by H-NS along DNA as a consequence of interactions between adjacent H-NS dimers (see above). The absence of cooperativity observed in the presence of Hha, indicates that Hha (unlike YdgT) interferes with filament formation. A higher effective affinity of H-NS-Hha to DNA compared to DNA bound H-NS-Hha may favor nucleation over filament formation (figure 3 - figure supplement 2b). The disparate effects of Hha and YdgT observed in relation to H-NS binding along DNA and comparable effects in relation to H-NS DNA-bridging, suggest that these proteins may drive a conformational switch (as seen for Mg^2+^) in addition to an altered DNA interaction surface. To investigate whether Hha affects H-NS conformation we performed MD simulations, incorporating structural information from the recently resolved H-NS_1-43_-Hha co-crystal structure^47^. Our MD simulations reveal that Hha does not prevent buckles in helix α3 (see figure 2b). Instead Hha alters the interactions between the dimerization domain and the DNA binding domain (Figure 2 - figure supplement 1) by blocking access to parts of dimerization domain. This hypothesis is further supported by interactions between Hha and other parts of H-NS, including the DNA binding domain and helix α3 (Figure 2 - figure supplement 7). In the presence of Mg^2+^ and Hha the contacts between the DNA binding domain and dimerization domain are reduced even further. This shows that Hha modulates H-NS function by stabilizing the “open” -bridging capable-conformation of H-NS.

## DISCUSSION

It has been known for many years that H-NS binding induces gene silencing. H-NS activity *in vivo* is modulated by physico-chemical factors (osmolarity, pH, temperature) and interaction partners. These findings support the hypothesis that H-NS plays a role in environmental adaptation. However, mechanistically it is unclear how functional modulation of H-NS by such factors is achieved. Based on our findings, we conclude that H-NS is incapable of bridging or stiffening DNA as dimers. H-NS dimers bind DNA *in cis* ^26,27^ and associate side-by-side along DNA, likely via the recently identified dimer-dimer interaction domain^24^, resulting in DNA stiffening. This process is cooperative as H-NS dimers interact with neighbors, as well as with DNA. Our studies reveal that H-NS-DNA filaments are *structurally* very similar, independent of the presence of Mg^2+^ (figure 1d). But *functionally*, these H-NS-DNA filaments are distinct. H-NS can be ‘activated' by Mg^2+^, which promotes a conformational change, rendering both DNA binding domains of H-NS dimers accessible for DNA bridging. In our model the assembly of bridged complexes proceeds in distinct steps: 1) nucleation ^48^ (binding of an H-NS dimer at a high affinity site), 2) lateral filament growth by H-NS dimer-dimer interactions (leading to DNA stiffening) and 3) bridging of the assembled filament to bare DNA provided *in trans* (figure 4). Each step can potentially be modulated by osmolarity and protein interaction partners. Here we show that these factors most effectively target DNA bridging.

**Figure 4.**
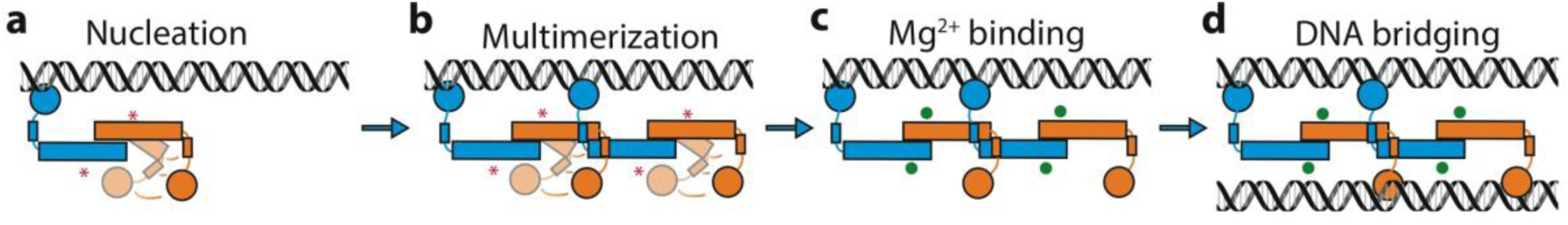
Model of H-NS complex assembly. (a) H-NS nucleates at preferred DNA sequences in the genome. (b) H-NS laterally multimerizes laterally along the DNA in the “closed” conformation. (c) In the presence of Mg^2+^ or other H-NS modilators such as Hha, H-NS switches to the “open”, bridging capable conformation. (d) H-NS forms DNA bridges *in trans.* The red asterisk indicates the buckle location. Mg^2+^ ions are shown as green orbs.

What are the implications of our observations? Our observations add to the large body of evidence showing that regulation of transcription via H-NS is complex, and that it does not proceed via a single, simple, well-defined mechanism. The most straightforward form of repression by H-NS is via occlusion of RNA polymerase from the promoter^49,50^. Whether this mechanism of repression involves lateral filament formation or bridging is unclear. It is expected that both types of complexes assembled at a promoter site can in principle occlude RNA polymerase. A second form of repression by H-NS is to interfere with RNA polymerase progression during active transcription by intragenic binding^51^. In this model both modes could interfere with transcription. However, it was suggested recently that only bridged filaments are capable of interfering with transcription^52^, with lateral filaments likely being disassembled after RNA polymerase encounters them. Generally, the type of complex formed by H-NS is expected to depend on the type and amount of ions, combined with local DNA conformation (dependent on DNA sequence or DNA topology), and the presence of modulating proteins. The interplay between these factors will determine the strength of the complex, and degree of repression. Thus, different H-NS repressed genes are expected to be subject to different types of modulation, providing a key to a coordinated response in gene expression to altered conditions and selectivity for the interplay with specific co-regulators at specific target regions.

## MATERIAL AND METHODS

### Construction of expression vectors

#### H-NS expression vector

The vector pRD18 for expression of H-NS was constructed by inserting the PCR amplified *hns* gene into the pET3His overexpression vector using NdeI and XhoI restriction sites. Using the XhoI restriction site the encoded protein does not contain a C-terminal His-tag.

#### H-NS_Y61DM64D_ expression vector

The vector pRD69 for expression of H-NS_Y61DM64D_ was constructed by inserting a PCR fragment containing the *hns* gene mutated to encode aspartic acid instead of tyrosine/methionine at position 61 and 64 into pET3His using NdeI and XhoI restriction sites.

#### Hha expression vector

The vector pRD38 for expression of N-terminally His tagged Hha was constructed by inserting a PCR fragment containing the *hha* gene into pET3His using XhoI and BamHI restriction sites.

#### YdgT expression vector

The vector pRD39 for expression of N-terminally His tagged YdgT was constructed by inserting a PCR fragment containing the *ydgT* gene into pET3His using XhoI and BamHI restriction sites.

### Protein overproduction and purification

#### H-NS/H-NS_Y61DM64D_

BL21 (DE3) Δ*hns*::*kan*/frt pLysE (NT201, our lab) cells transformed with plasmids expressing H-NS/H-NS mutants were grown to an OD_600_ of 0.4, and induced for two hours using IPTG (500 μM). The cells were pelleted and lysed by sonication in 100 mM NH_4_Cl, 20 mM Tris pH 7.2, 10% glycerol, 8 mM β-mercaptoethanol, 3 mM benzamidine). The soluble fraction was loaded onto a P11 column and eluted using a 100 mM-1 M NH_4_Cl gradient, the protein eluted at 280 mM NH_4_Cl. The peak fractions were dialysed to buffer B (identical to buffer A, but containing 130 mM NaCl instead of NH_4_Cl) by overnight dialysis. The dialysate was loaded onto a heparin column (GE Healthcare) and eluted using a 130 mM-1 M NaCl gradient, the protein eluted at 350 mM NaCl. The pooled peak fractions were dialysed to buffer B and concentrated using a 1 ml Resource-Q column (GE Healthcare). The purity of the protein was verified on an SDS-PAGE gel. The protein concentration was determined using a Bicinchoninic Acid assay (Pierce BCA protein assay kit, Thermo Scientific).

#### H-NS_1-58_

BL21 (DE3) Δ*hns*::*kan*/frt pLysE (NT210, our lab) cells transformed with plasmids expressing H-NS/H-NS mutants were grown to an OD_600_ of 0.4, and induced for two hours using IPTG (500 μM). The cells were pelleted and lysed by sonication in 100 mM NaCl, 20 mM Tris pH 7.2, 10% glycerol, 8 mM β-mercaptoethanol, 3 mM benzamidine). The soluble fraction was heated to 65°C for 10 minutes and then spun down at 10.000 RPM for 10 minutes. The supernatant was collected and a 1:1 ratio saturated ammonium sulfate (50 mM Tris pH 7.2, 4M ammonium sulfate) was gradually added to the cooled sample. The sample was spun down at 8.000 RPM for 15 minutes and a 1:1 ratio of 5 mM Tris pH 7.2, 15% glycerol was added to the supernatant. To remove further impurities the sample was run through a 1ml hydrophobic interaction column and 1ml Blue-agarose column (the protein should not bind to either of these column) before finally binding the protein to 1 ml Resource-Q column (GE Healthcare). The protein was eluted with a 25mM -1M gradient of NaCl, the protein eluted at roughly 380 mM NaCl. The purity of the protein was verified on an SDS-PAGE gel. The protein concentration was determined using a Bicinchoninic Acid assay (Pierce BCA protein assay kit, Thermo Scientific).

#### Hha/YdgT

BL21 (DE3) Δ*hns*::frt, *hha*::*kan*, pLysE (our lab) cells transformed with plasmids pRD38/pRD39 expressing *hha/ydgT* were grown at 37°C to an OD_600_ of 0.4, and induced for two hours using IPTG (500 μM). The cells were pelleted and lysed in 20 mM HEPES pH 7.9, 1 M KCl, 10 % glycerol, 8 mM β-mercaptoethanol. The soluble fraction was loaded onto a Ni-column. The column was first washed with buffer D (20 mM HEPES pH 7.9, 0.5 M KCl, 10% glycerol, 8 mM β-mercaptoethanol). The protein was then eluted using a 0 mM-0.5 M Imidazole gradient, the protein eluted at 300 mM Imidazole. The peak fractions were dialysed to buffer E (identical to buffer D, but containing 100 mM KCl) by overnight dialysis. The sample was then loaded onto pre-equilibrated SP Hi-Trap-column and Ni-column connected in series. After loading the samples on the column, the SP Hi-Trap -column was disconnected and the protein was eluted from the Ni-column using a 0-0.5 M imidazole gradient. The purity of the protein was verified on an SDS-PAGE gel. The protein concentration was determined using a Bicinchoninic Acid assay.

### Peptide production and purification

A truncated form of H-NS (H-NS_56-82_) was synthesized by way of automated solid phase synthesis was performed using standard protocols via Fmoc-strategy. Purification was performed by RP-HPLC with a Gemini 5μ C18 reversed phase column. Identity of the peptides was determined via MALDI-MS. The purity was determined by means of analytical RP-HPLC. The peptide was freeze dried and dissolved in 20 mM Tris pH 7.2, 300 mM KCl, 10% glycerol, 8 mM β-mercaptoethanol. The peptide concentration was determined using a Bicinchoninic Acid assay (Pierce BCA protein assay kit, Thermo Scientific).

## Size exclusion chromatography

Size exclusion chromatography was done using a Superose-12 column with a flow of 0.3 ml/min, pre-equilibrated with 10 mM Tris-HCl, 50 mM KCl, 5% glycerol containing or lacking 10 mM MgCl_2_. The absorbance of the eluting fractions was measured at 215 nm. These experiments were performed in triplicate.

## DNA substrates

### DNA preparation

All experiments were performed using a random, AT rich, 685 bp (32% GC) DNA substrate described in literature^53^. The DNA substrate was generated by PCR, and the PCR products were purified using a GenElute PCR Clean-up kit (Sigma-Aldrich). If required, DNA was ^32^P-labeled as described previously ^54^.

## DNA bridging assay

Streptavidin-coated paramagnetic Dynabeads M280 (Invitrogen) were washed once with 100 μL of 1xPBS and twice with Coupling Buffer (CB: 20 mM Tris-HCl pH 8.0, 2 mM EDTA, 2 M NaCl, 2 mg/mL BSA(ac), 0.04% Tween20) according to manufacturer instructions. After washing, the beads were resuspended in 200 μL CB containing 100 nM biotinylated DNA. Next, the bead suspensions were incubated for 30 minutes on a rotary shaker (1000 rpm) at 25°C. After incubation, the beads were washed twice with Incubation buffer (IB: 10 mM Tris-HCl pH 8.0, 50 mM KCl, 10* mM MgCl_2_, 5% v/v Glycerol, 1 mM DTT and 1 mM Spermidine) before resuspension in IB and addition of ±8000 cpm of radioactively labeled ^32^P 685 bp DNA. Radioactive DNA was supplemented with unlabeled 685 bp DNA to maintain a constant (20 nM) DNA concentration. The DNA bridging protein H-NS (concentrations indicated in the text), and if applicable Hha or YdgT were added and the mixture was incubated for 30 minutes on a shaker (1000 rpm) at 25°C. To remove unbridged prey DNA, the beads were washed with IB, before resuspension in 12 μL stop buffer (10 mM Tris pH 8.0, 1 mM EDTA, 200 mM NaCl, 0.2% SDS). All samples were quantified through liquid scintillation counting over 10 minutes. All values recovered from the DNA bridging assay were corrected for background signal (using a sample lacking H-NS), and normalized to a reference sample containing the amount of labeled ^32^P 685 bp DNA used in the assay. The samples were then run on a 5% 0.5x TBE gel to ensure DNA integrity. DNA bridging was calculated based on a reference sample containing 2 μL of prey DNA. All DNA bridging experiments were performed in triplicate. Unless indicated otherwise all DNA bridging experiments were performed in the presence of 10 mM of MgCl_2_.

## Tethered particle motion experiments

Tethered particle motion experiments were performed as reported previously^55,56^. Flow cells were prepared as described with minor modifications^55,56^. Here, before flowing in protein diluted in the experimental buffer (10 mM Tris-HCl pH 8.0, 50 mM KCl, 10 mM MgCl_2_/EDTA, 5% v/v Glycerol, 1 mM DTT) the flow cell was washed using 4 flow cell volumes with the experimental buffer. Next, the flow cell was incubated for 10 minutes with protein solution before sealing the flow cell. The flow cell was maintained at a constant temperature of 25 °C. Measurements were started 10 min. after the introduction of protein solution. TPM experiments were done at least in duplicate. The data were analyzed as previously described^55,56^. The fractional coverage was fit using the McGhee-von Hippel model for cooperative lattice binding ^57^. To this end, weighted orthogonal distance regression (ODR) was performed to estimate the parameters of the nonlinear implicit equation describing the cooperative ligand binding. The binding site size (n) of H-NS was fixed to a value of 30 bp during regression, which corresponds to values determined previously^26,31^. The association constant (K) and the cooperativity parameter (ω) were assumed to be positive real numbers. A custom fitting routine was implemented in Fortran and makes use of the ODRPACK library ^58^.

## Molecular Dynamics simulations

The starting conformation of the full length H-NS dimer was constructed as described previously^14^. The system was placed in a periodic dodecahedron box with a distance of at least 0.8 nm between the box edge and the most extended atom of the protein dimer, followed by the addition of water and ions. With this system we performed Molecular Dynamics (MD) simulations of full length H-Ns at different concentrations of KCl and MgCl_2_ and with the addition of Hha, summing up to a total of six unique systems. Hha was added by aligning the crystal structure containing the H-NS – Hha complex (PDB code 4ICG ^47^) with the full-length structural model and copying the Hha molecules. System size ranged from 513457 atoms for the Hha systems to around 1135000 atoms for the other four systems.

Interactions between atoms were described by the AMBER99-SB-ILDN force field ^59^, in combination with the TIP3P water model ^60^. Long-range electrostatic interactions were treated via the Particle Mesh Ewald method ^61,62^ with a short-range electrostatic cutoff distance at 1.1 nm. Van der Waals interactions were cut off at 1.1 nm. Preparation of the systems consisted of energy minimization equilibration of the solvent. Energy minimization was performed by the conjugate gradient method. After energy minimization, the positions of water molecules and ions were equilibrated by a 1 ns molecular dynamics run at a temperature of 298 K and a pressure of 1 bar in which the heavy atoms in the protein were position-restrained with a force constant in each direction of 1000 kJ/mol nm. After preparation we performed 16 50 ns runs for each system, varying initial conditions by assigning new random starting velocities drawn from the Maxwell-Boltzmann distribution at 298 K. All simulations were performed with GROMACS v.4.6.3 ^63^ at the Dutch National Supercomputer with the leapfrog integration scheme and a time step of 2 fs, using LINCS ^64^ to constrain all bonds. All simulations were performed in the isothermal-isobaric ensemble at a pressure of 1 bar, using the v-rescale thermostat ^65^ and the Parrinello-Rahman barostat ^66^.

Frames were stored every 10 ps. The first 10 ns of each simulation are excluded from analysis, unless stated otherwise. Analysis focused on determining contacts between domains, between H-NS and Hha, between H-NS and ions, and helical hydrogen bonds. Contact maps of interactions between residues in the H-NS dimer system were obtained by first calculating the minimum distance between each residue pair in the system. A residue pair is counted to be in contact if they are at a minimum distance of 0.6 nm or less. The probability of a contact is then calculated as the average over all 16 simulations (excluding the first 10 ns) and displayed as a contact probability matrix. We used a modified version of the g_mdmat tool in GROMACS^63^ in combination with Perl scripts to generate contact maps. To determine the location of magnesium with respect to the H-NS system, we calculated the minimum distance between each residue in the H-NS dimer and the Mg^2+^ ions and counted a contact if the distance between an H-NS residue and a magnesium ion is 0.6 nm or less. These contact probabilities P_Mg2+_are averaged over all 16 simulations and the two monomers. A similar procedure was performed to determine the probability of contacts between H-NS and Hha, PHha, and between H-NS domains, with P_1-40_ indicating the probability of finding the DNA binding domain (residues 96-137) close to the dimerization domain (residues 1-40). P_96-137_ indicates the probability of finding the dimerization domain close to the DNA binding domain.

To determine the location of the buckle in helix α3, we calculated the helical hydrogen bond distances d_O-N_ for residues 22-67 in each monomer between the backbone carbonyl oxygen O of residue i and the backbone amide nitrogen N of residue i+4. A hydrogen bond is counted to be in contact if they are at a minimum distance of 0.35 nm or less. These probabilities are averaged over all 16 simulations (excluding the first 10 ns) and the two monomers. Snapshots and movies were generated with PyMOL.

## ACKNOWLEDGMENTS

We thank Bas de Mooij for his assistance in standardizing the bridging assay. We also thank Wim Jesse and Alexander Kros for synthesizing the H-NS56-83 peptide and their assistance during its purification. We also thank Rosalie Driessen for her assistance with data analysis and valued discussions. JV acknowledges the use of the Dutch National Supercomputer Cartesius for the MD simulations.

## Additional Information

The authors declare no competing financial interests.

## Supplementary Material

**Figure 1 - figure supplement 1:**
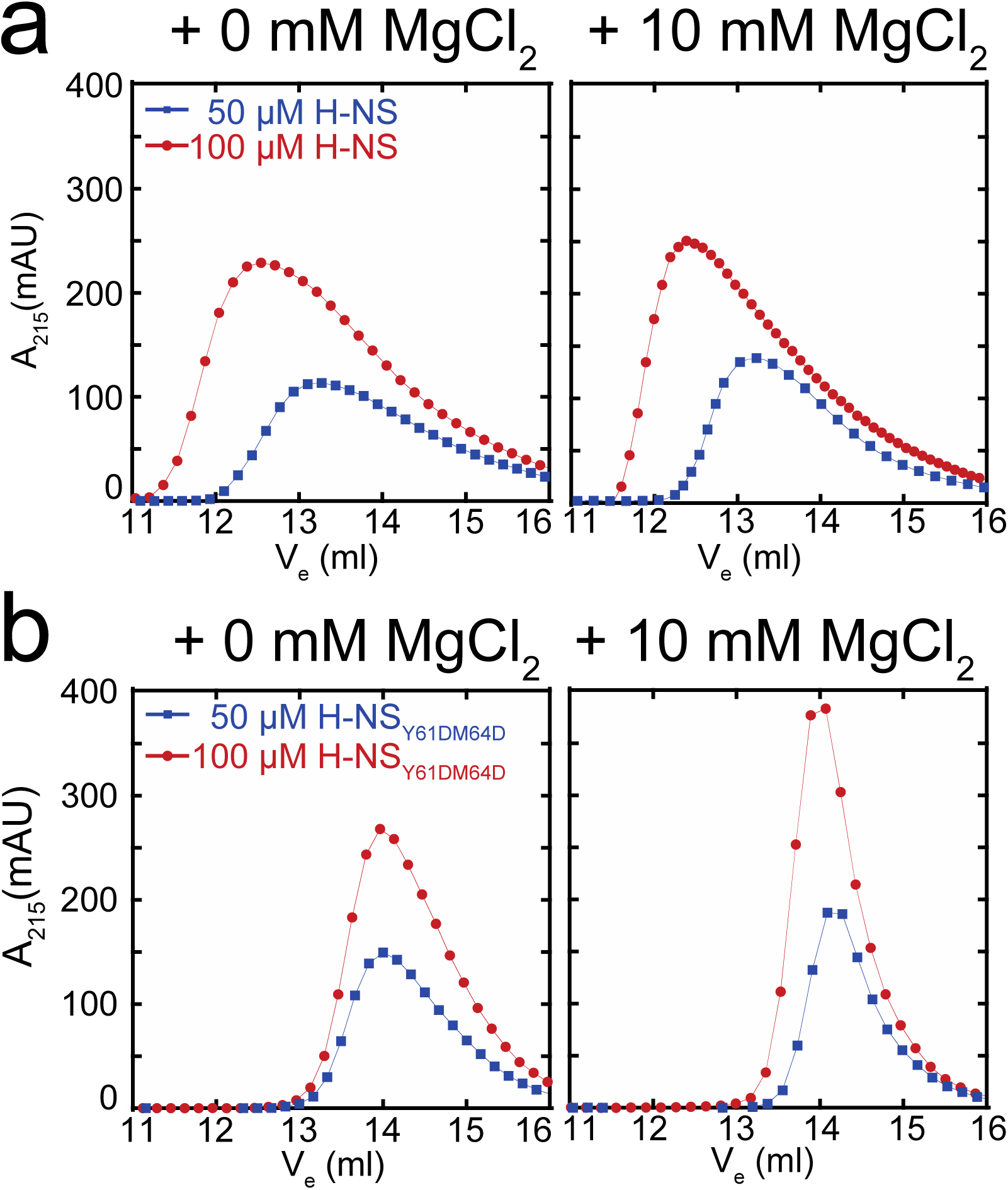
Multimeric state of H-NS measured using size exclusion chromatography. (a) Elution pattern of 50 and 100 μM of H-NS as a function of Mg^2+^. (b) Elution pattern of 50 and 100 μM of H-N_SY61DM64D_ as a function of Mg^2+^. Lines are to guide the eye.

**Figure 1 - figure supplement 2:**
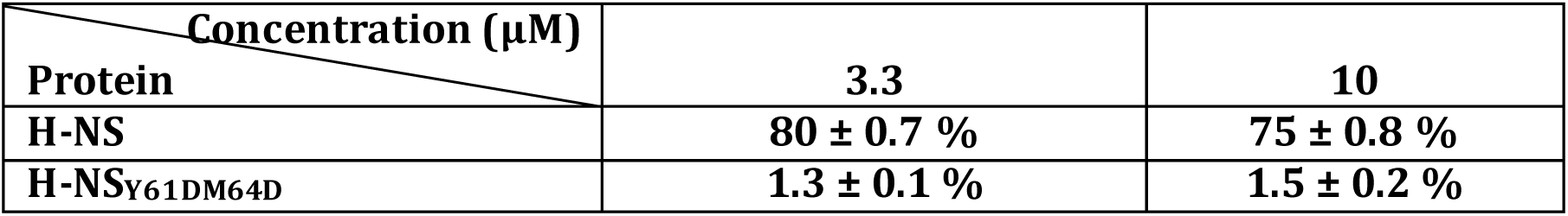
DNA bridging efficiency of H-NS and H-NS_Y61DM64D_.

**Figure 1 - Supplemental Figure 3:**
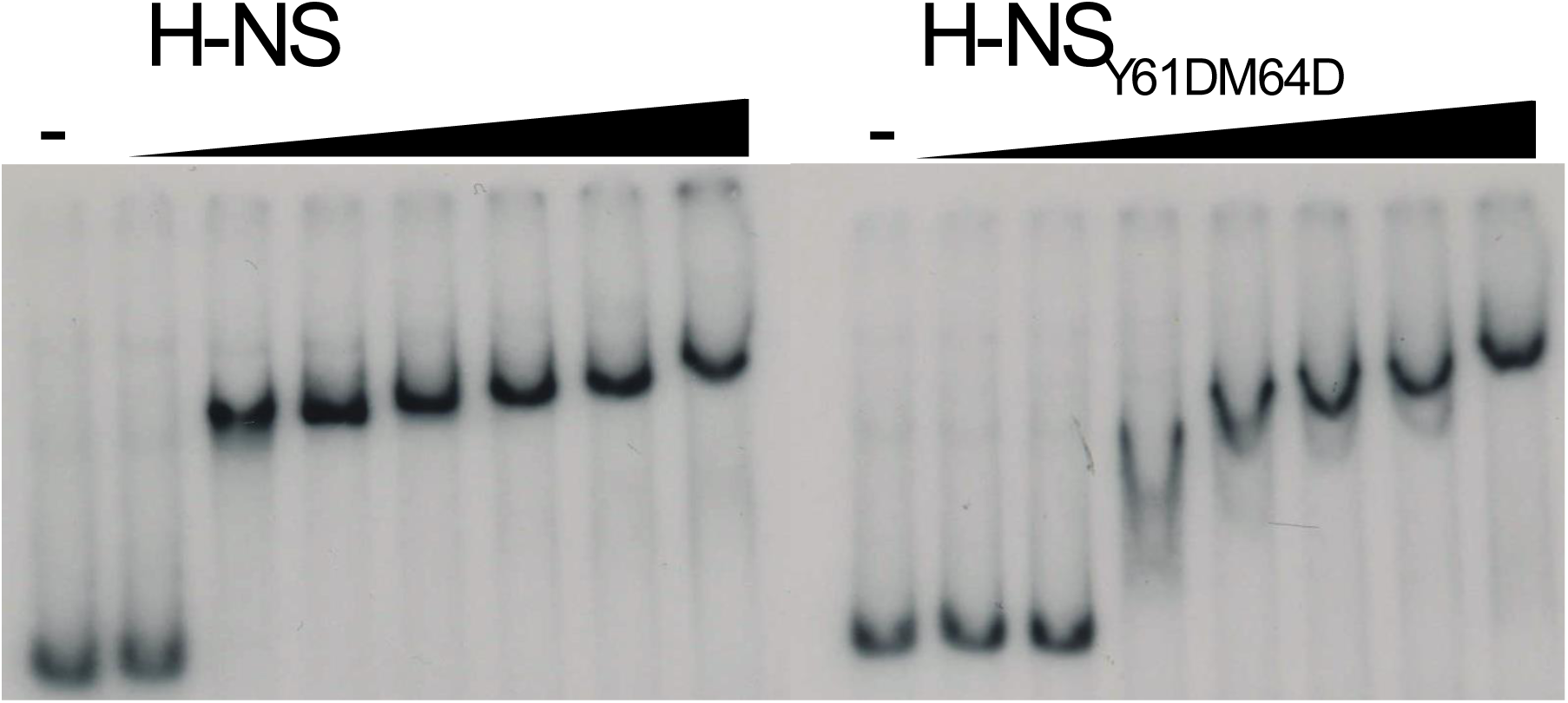
Electrophoretic Mobility shift assay. (a) H-NS and (b) H-NS_Y61DM64D_. The proteins were added to ^32^P-labeled 685 bp DNA substrate at 0.4,0.8,1.2,1.6,12.4,3.3,6.6 μM concentrations. Both H-NS and H-NST_Y61DM64D_ have very similar DNA binding affinities with approximate Kd values of 1 ± 0.2 μM for H-NS and 1.4± 0.2 μM for H-NS_Y61DM64D_ respectively. This small difference can in part be attributed to a difference in cooperativity of DNA binding between the two variants on H-NS.

**Figure 1 - Supplemental Figure 4:**
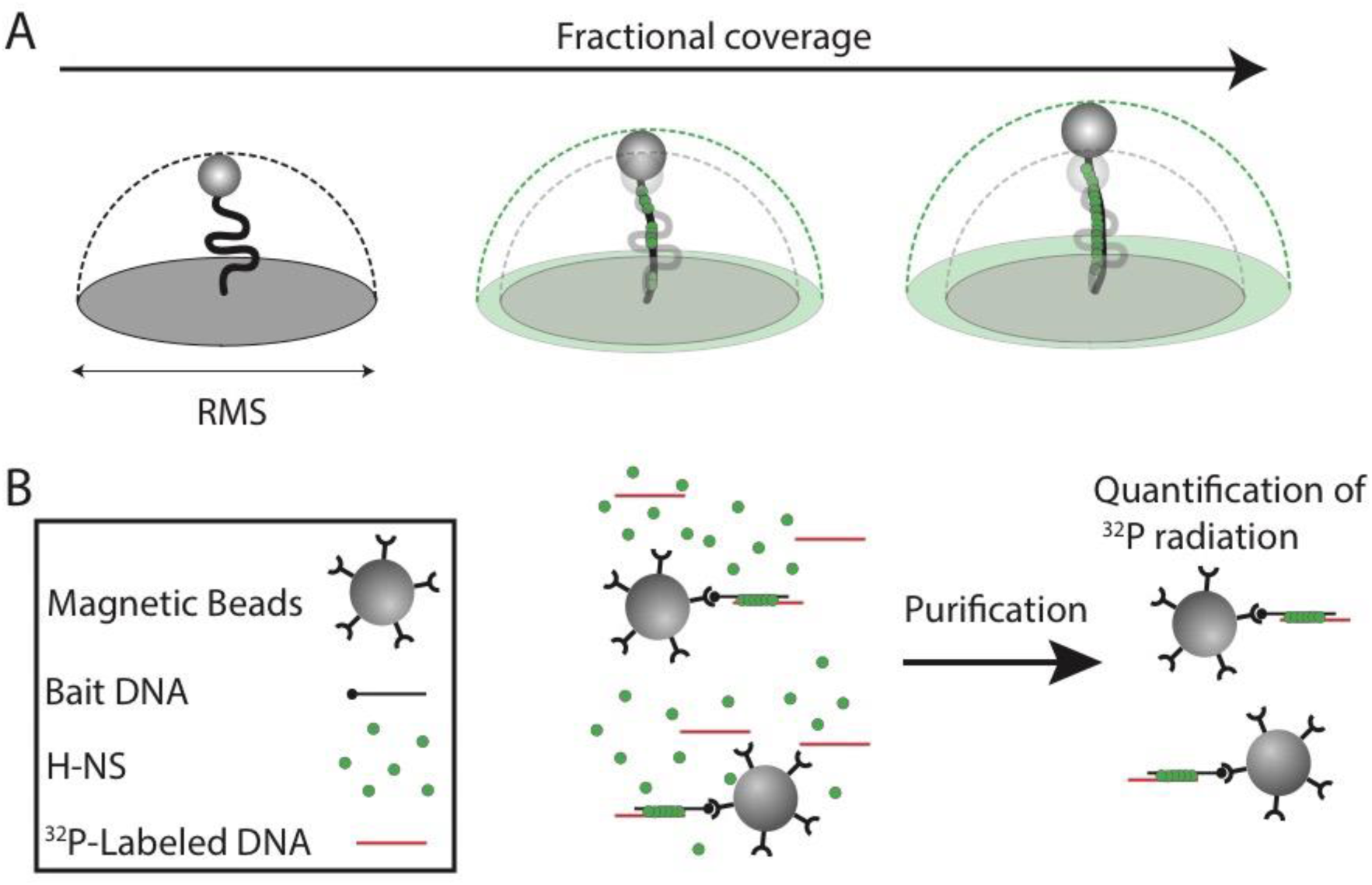
Schematic depiction of techniques used in this study. (a) Schematic depiction of TPM and the effect of increased protein binding to DNA on the Root Mean Squared (RMS) of the bead. (b) Schematic depiction of the DNA bridging assay, where a streptavidin coated paramagnetic bead is coupled to a 5’ biotin labeled bait DNA. These are then incubated in the presence of ^32^P radiolabeled DNA strand and H-NS. The paramagnetic beads are then pulled down and the amount of recovered prey DNA is quantified.

**Figure 2 - figure supplement 1:**
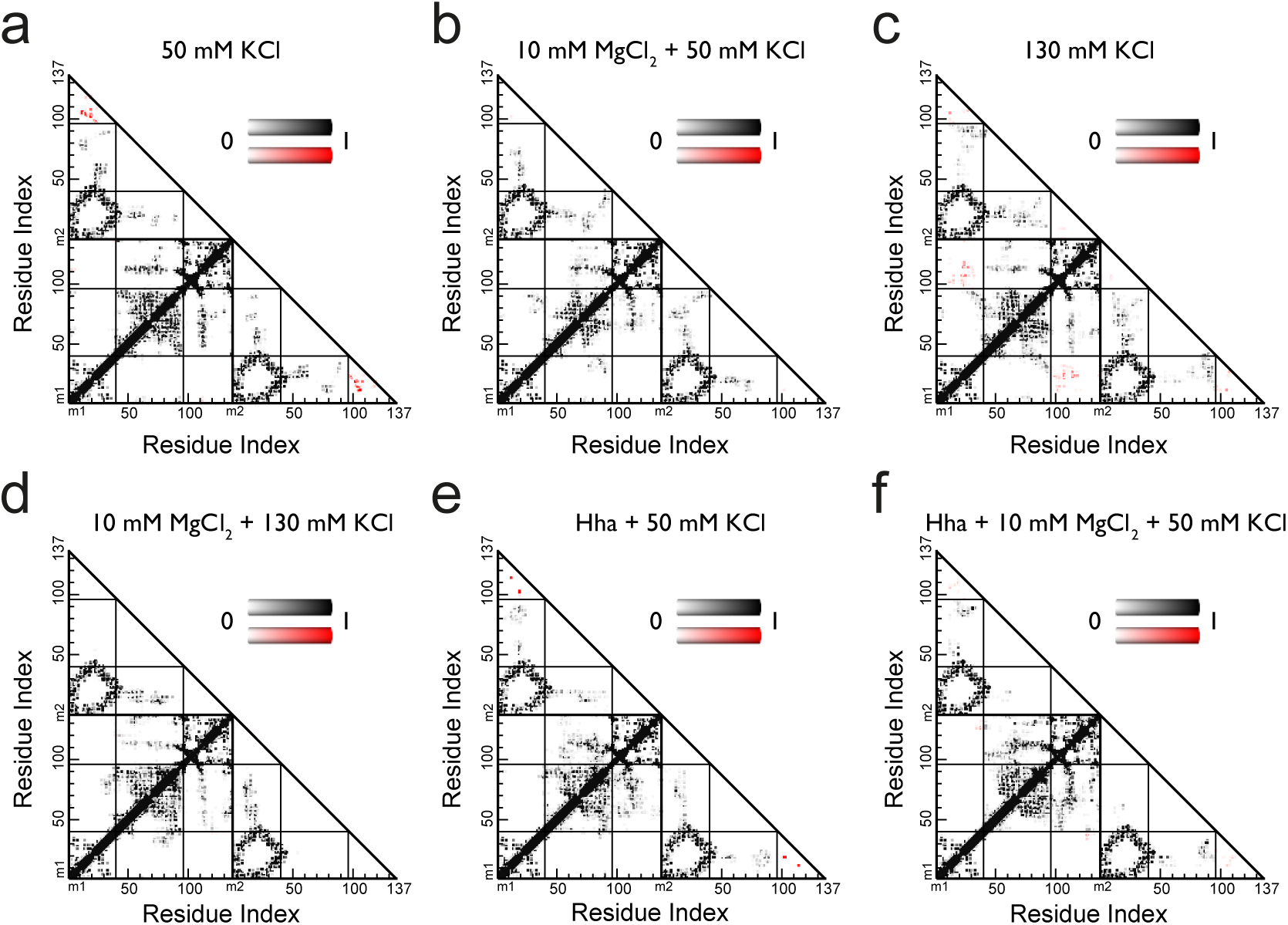
Contact maps of full-length H-NS dimers simulations in different conditions. (a) 50 mM NaCl, (b) 10 mM MgCl_2_ + 50 mM NaCl, (c) 130 mM NaCl, (d) 10 mM MgCl_2_ + 130 mM NaCl, (e) Hha + 50 mM NaCl, (f) Hha + 10 mM MgCl_2_ + 50 mM NaCl, Residues are considered in contact if the smallest distance between two atoms is 0.6 nm or less. The maps show the probability of finding a contact between residues, averaged over the last 40 ns of 8 molecular dynamics runs per system. White represents a probability of zero, darker colors indicates higher probability. The color scale runs from 0 to 1. Contacts between the dimerization domain and the DNA binding domain are colored red.

**Figure 2 - Supplemental Figure 2: Conformational flexibility of H-NS.** The protein is rendered in cartoon representation with a transparent surface. The color code indicates the domain organization of H-NS: blue - site 1 (residue 1-40), red - the buckle region in helix 3 (residue 42-49), green - site2 (residue 55-83) and brown - the DNA binding domain (residue 96-137). The DNA binding site on H-NS is shown in sticks. This movie contains snapshots at every 100 ps from run5 of the MD simulations of H-NS at 50 mM KCl (see Figure 2 - figure supplement 3 for time traces of the helical hydrogen bond distance between residue 45 and 49). The snapshots are rendered with PyMol.

**Figure 2 - Supplemental Figure 3:**
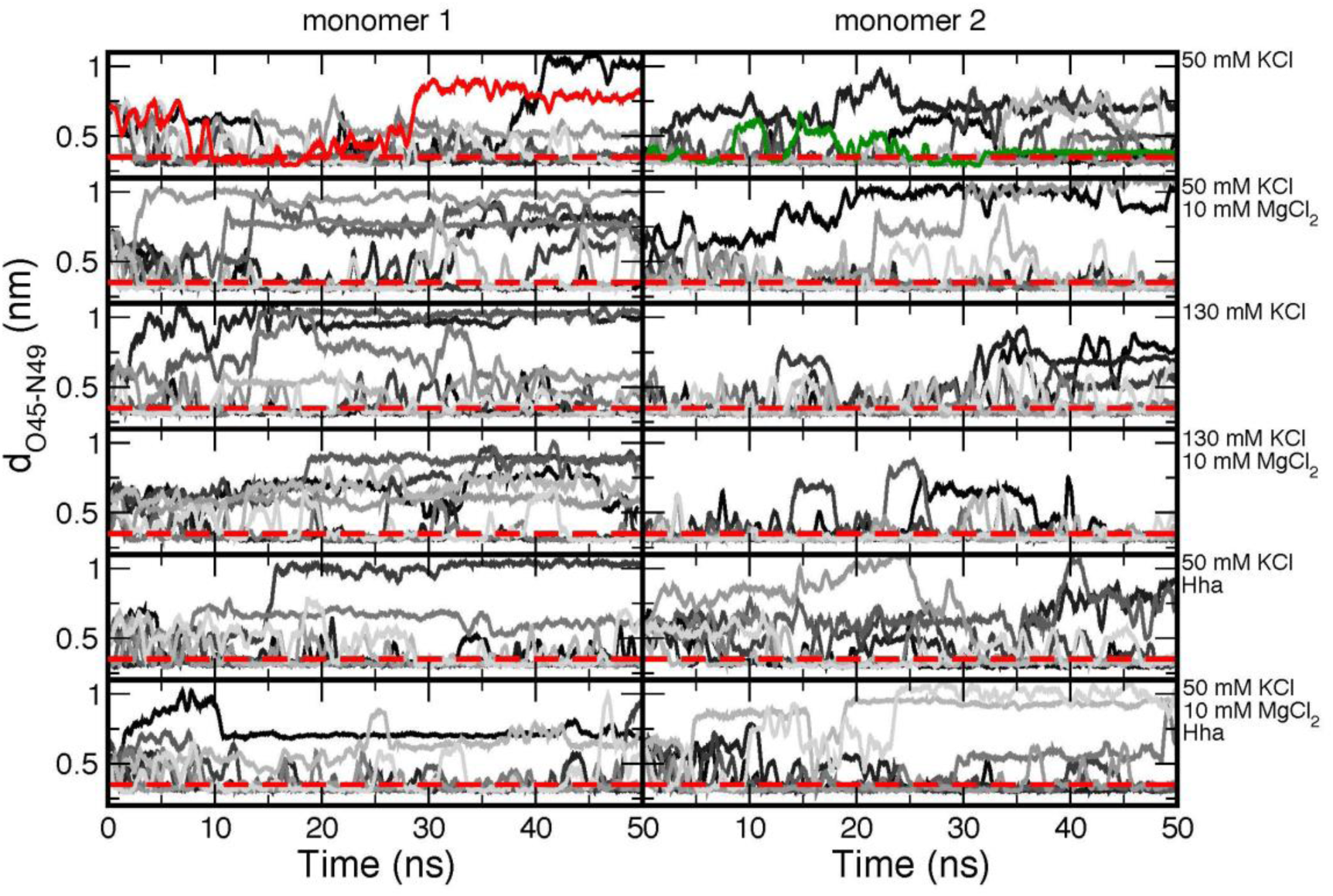
Time traces of the O-H distance between residues 45 and 49. The time traces are shown in different shades of gray to indicate the different runs. The red dashed line indicates the distance threshold for forming a hydrogen bond. The red and green lines are examples of the formation of irreversible and reversible buckles respectively. The time traces are shown as a running average of 50 ps to reduce the noise from thermal fluctuations.

Figure 2 - Supplemental Figure 4:The effect of magnesium on the conformational flexibility of H-NS. The protein is rendered in cartoon representation with a transparent surface. The color code indicates the domain organization of H-NS: blue - site 1 (residue 1-40), red - the buckle region in helix 3 (residue 42-49), green - site2 (residue 55-83) and brown - the DNA binding domain (residue 96-137). The DNA binding site on H-NS is shown in sticks. This movie contains snapshots at every 100 ps from run7 of the MD simulations of H-NS at 50 mM KCl + 10 mM MgCl_2_ (see Figure 2 - figure supplement 3 for time traces of the helical hydrogen bond distance between residue 45 and 49 and the minimum distance between residue 45 and the magnesium ions). The snapshots are rendered with PyMol.

**Figure 2 - Supplemental Figure 5:**
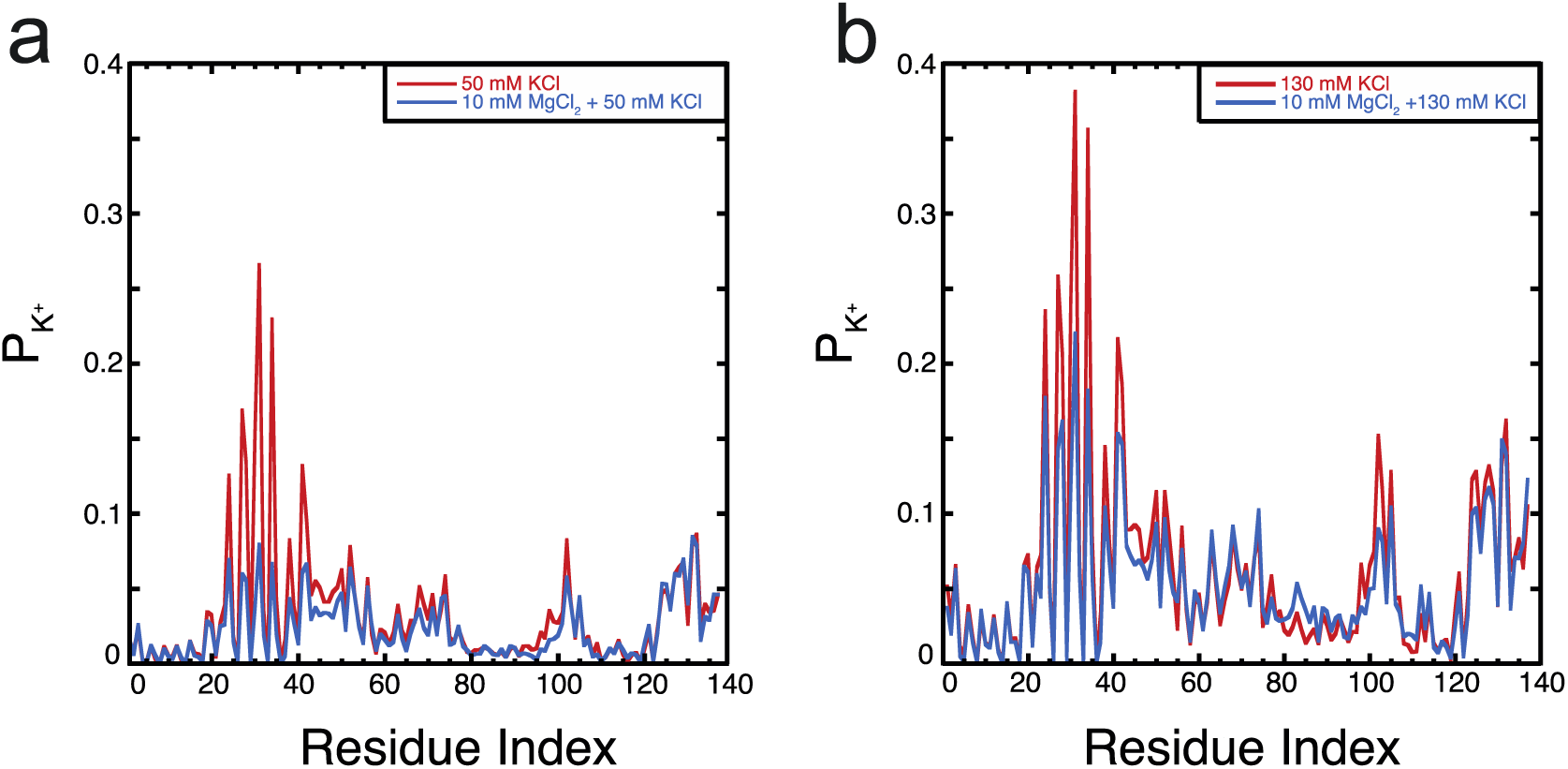
Location of K^+^ on H-NS. The probability of finding K^+^ ions within 0.6 nm of an H-NS residue, P_k+_, is plotted as function of the residue index for (a) 50 mM KCl and (b) 130 mM KCl in the presence and absence of MgCl_2_.

**Figure 2 - Supplemental Figure 6:**
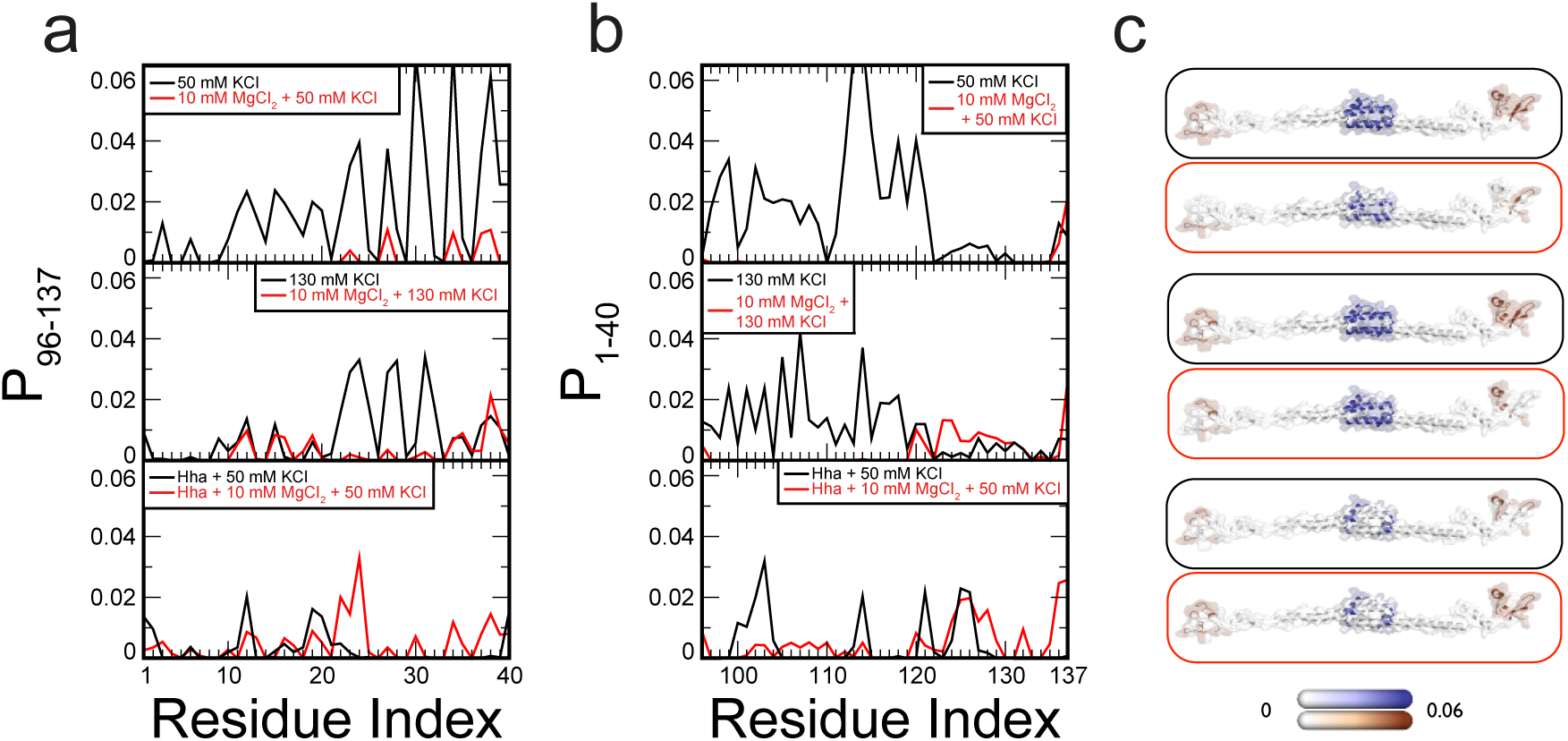
Contact maps of H-NS dimers in different conditions, focused on the interactions between the dimerization domain and the DNA binding domain. (a) The probability of finding the DNA binding domain within 0.6 nm of residues in the dimerization domain, P_96-137_ (b) The probability of finding the dimerization domain within 0.6 nm of residues in the DNA binding domain, P_1-40_. Residues are considered in contact if the smallest distance between two atoms is 0.6 nm or less. (c) P_96-137_ and P_1-40_ are highlighted on a three-dimensional structural model of H-NS. The protein dimer is shown in ribbon representation and transparent surface representation, with white indicating no interactions, blue indicating values for P_96-137_ and brown indicating P_96-137_. The color scale runs from 0 to 0.06.

**Figure 2 - Supplemental Figure 7:**
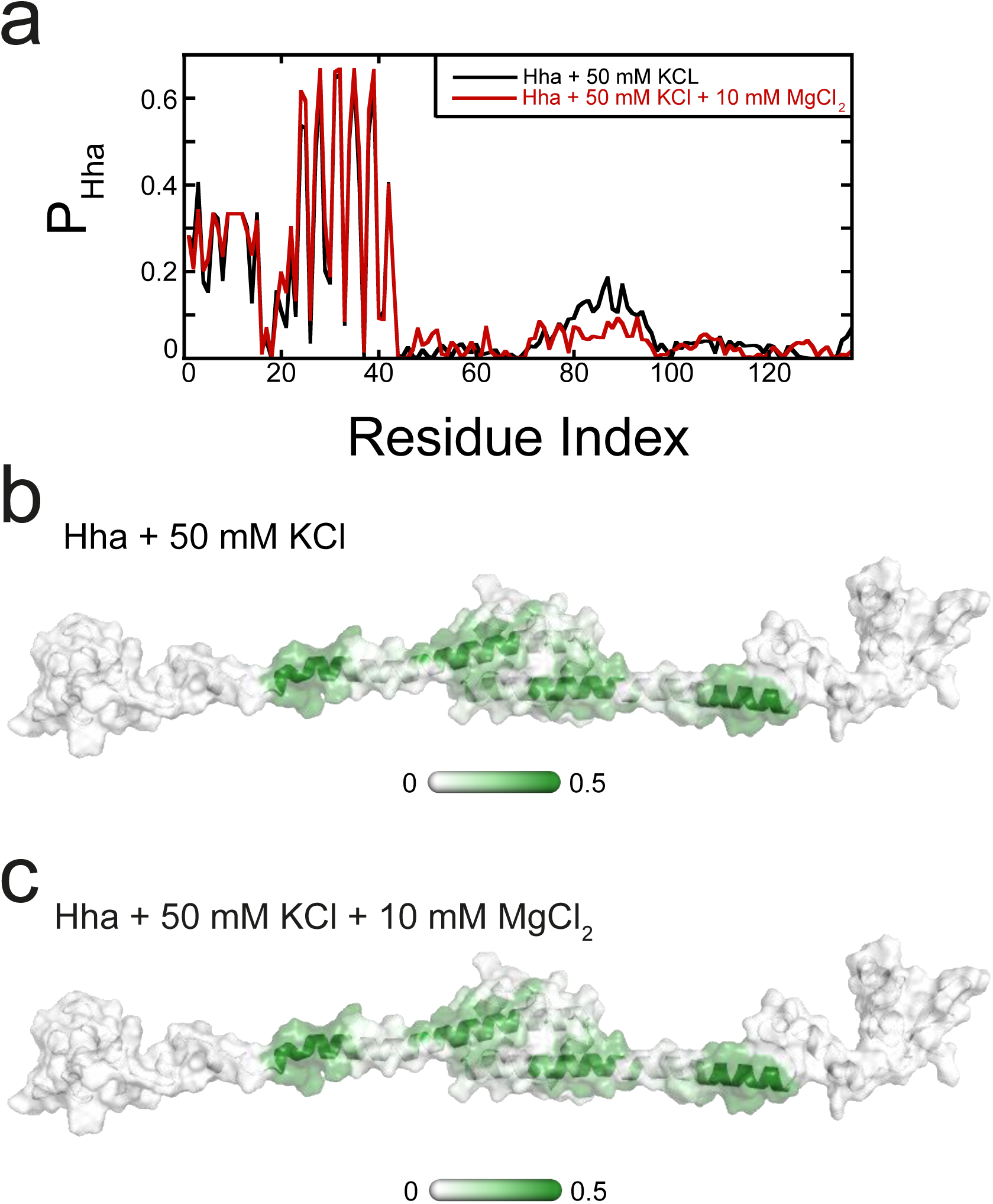
Location of Hha on H-NS. (a) The probability of finding Hha within 0.6 nm of an H-NS residue, P_Hha_, is plotted as a function of the residue index. The P_Hha_ is indicated on a surface representation of an H-NS dimer in open conformation in the (b) absence and (c) presence of MgCl_2_, ranging from 0 (white) to 0.5 (dark green).

**Figure 2 - Supplemental Figure 8:**
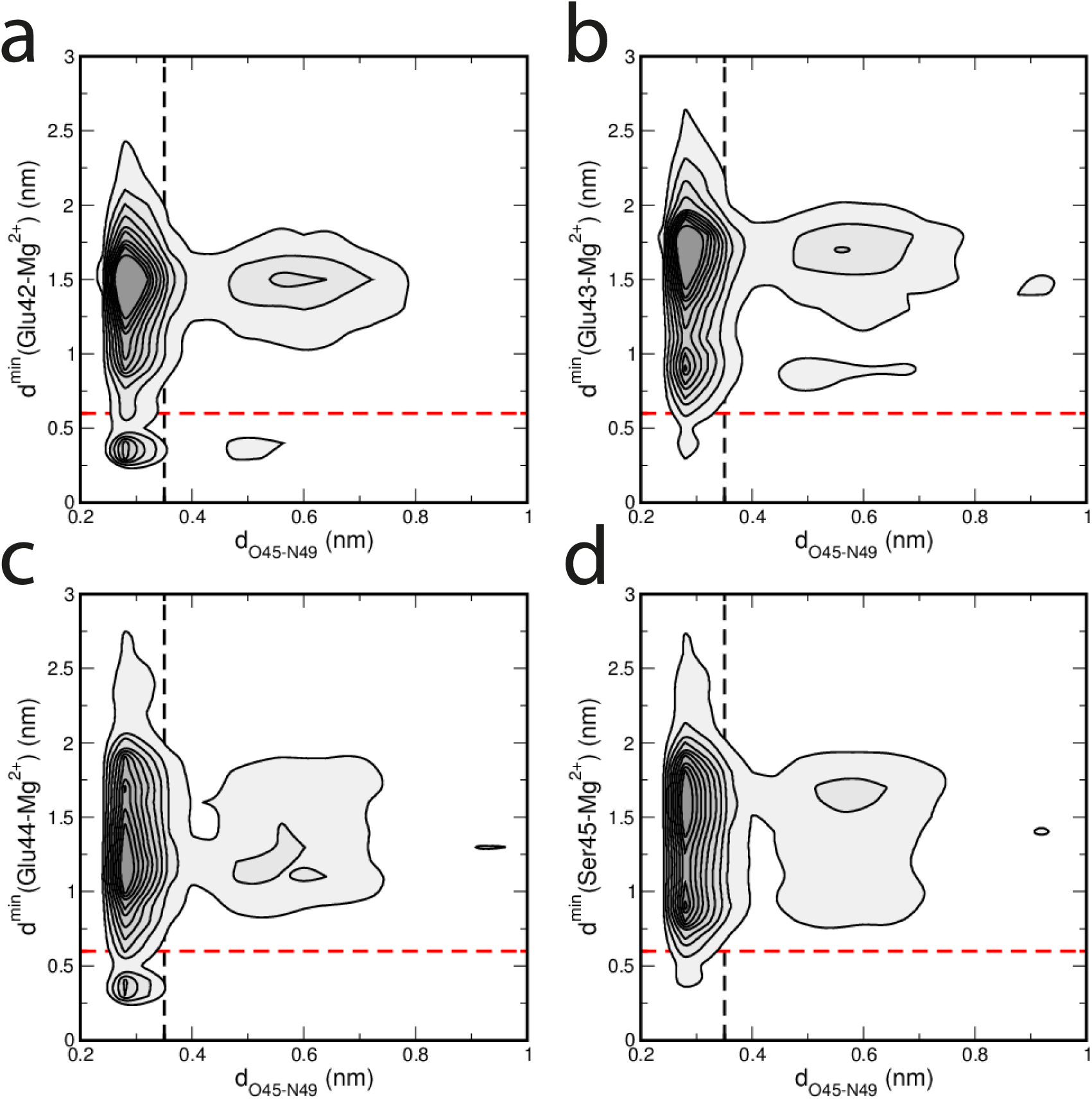
Correlation between hydrogen bond distance and proximity of Mg^2+^ to the buckle in helix α3. The correlation between the hydrogen bond distance d_O45-N49_ and the minimum distance between Mg^2+^ ions and a residue d^min^ is shown as a contour plot. The darker colours indicate higher probabilities of finding that particular configuration. Contours are drawn for every 0.2%. A distance larger than 0.35 nm indicates a broken hydrogen bond (black dashed line). A Mg^2+^ ion is close if this distance is less than 0.6 nm (red dashed line). The panels show the contour plots for four residues: (a) Glu42, (b) Glu43, (c) Glu44 and (d) Ser45.

**Figure 2 - Supplemental Figure 9:**
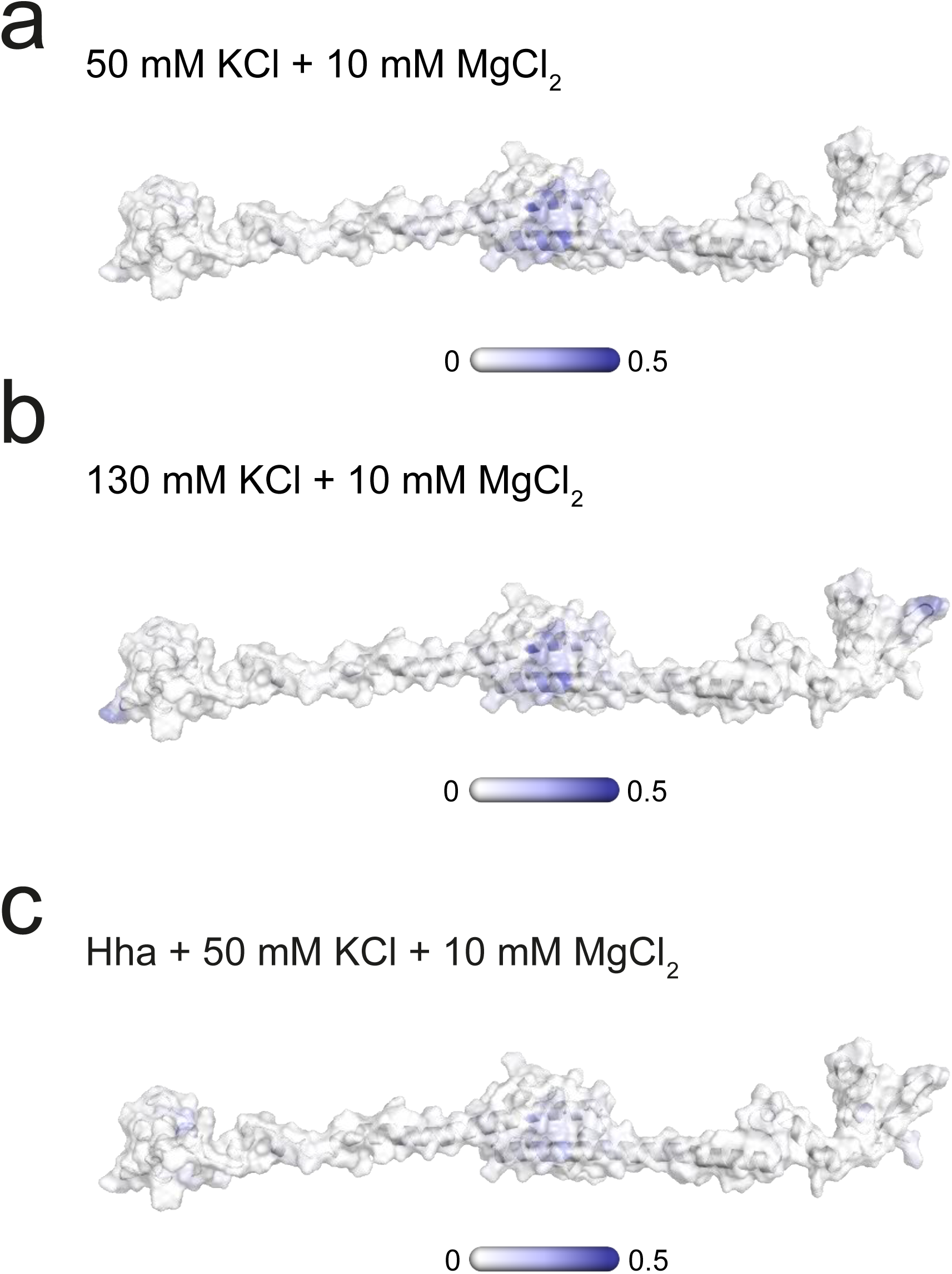
Mg localization on H-NS. The probability of finding Mg^2+^within 0.6 nm of H-NS in the presence of (a) 50 mM KCl, (b)130 mM KCl, or (c) Hha. This probability is indicated on a surface representation of an H-NS dimer in open conformation, ranging from 0 (white) to 0.5 (dark blue).

**Figure 3 - figure supplement 1:**
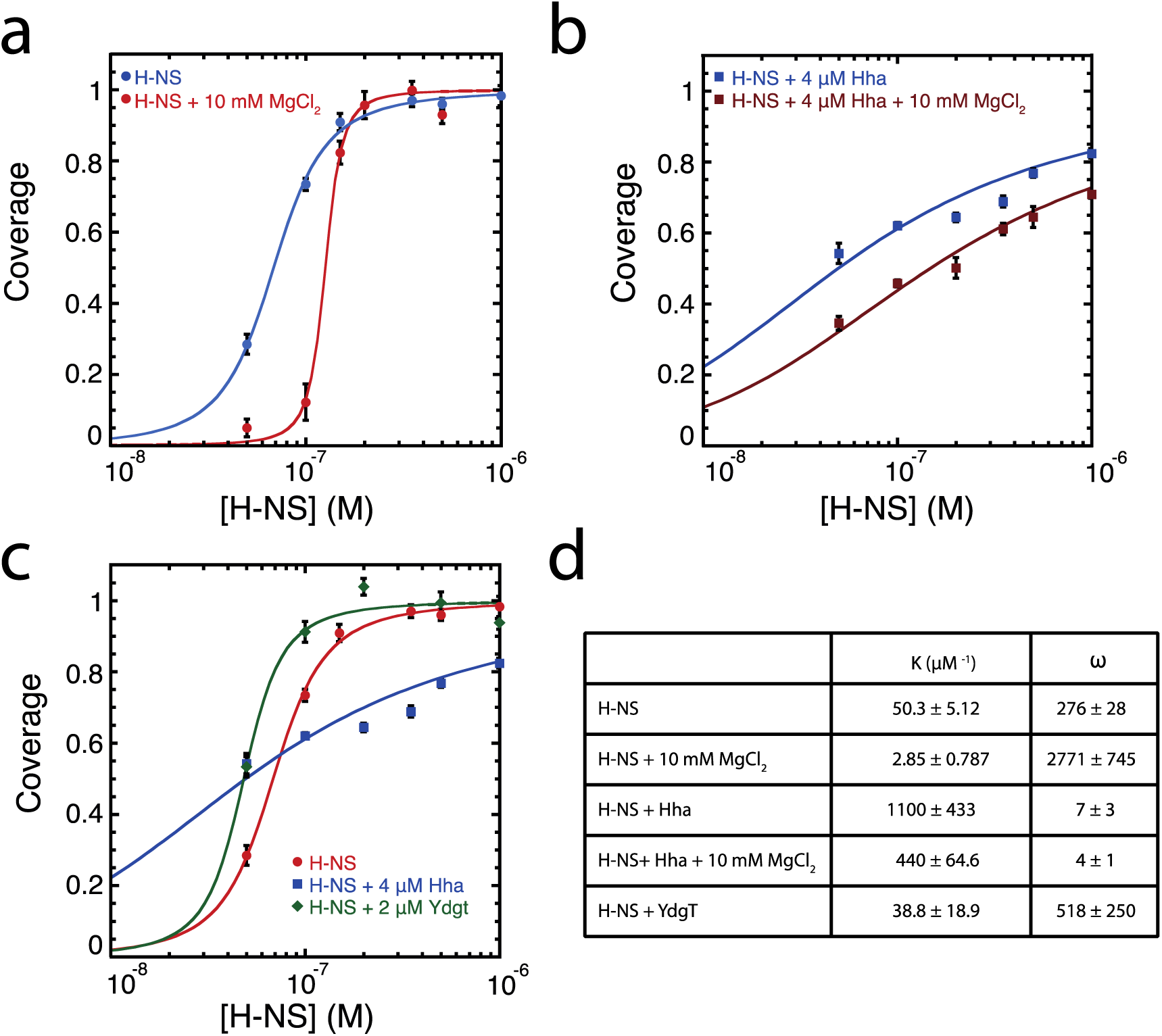
McGhee-von Hippel analysis of H-NS DNA binding curves based on TPM data. Binding curves of (a) H-NS, (b) H-NS + Hha TPM data as a function of Mg^2+^, and (c) H-NS, H-NS + Hha, and H-NS + YdgT. (d) Fit variables for all datasets, the binding affinity (K), and cooperativity (ω). Fit curves are shown as solid lines.

**Figure 3 - figure supplement 2:**
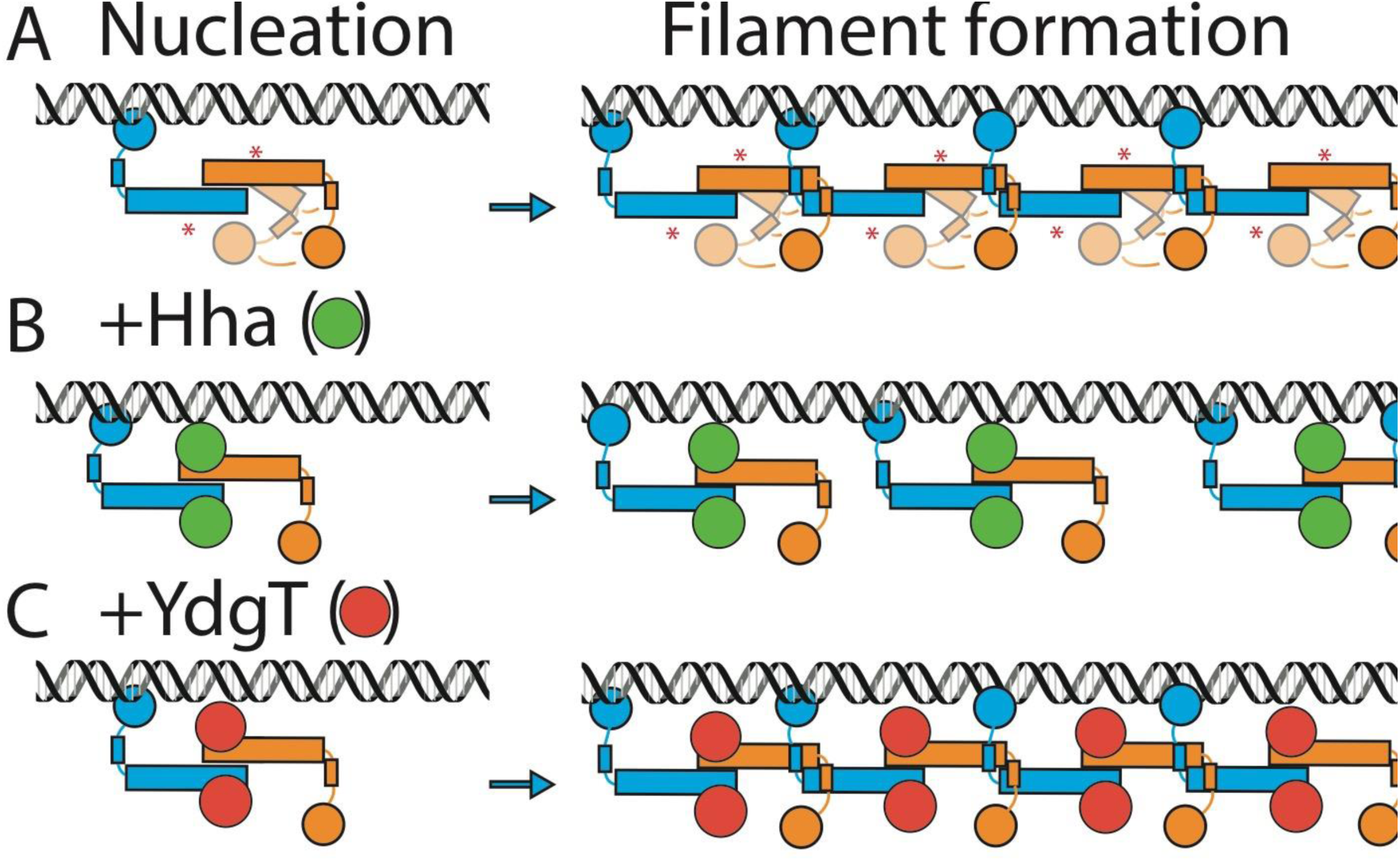
Schematic depiction of the effects of Hha and YdgT. Schematic depiction of (a) H-NS (b) H-NS + Hha, and (c) H-NS + YdgT, binding. H-NS or H-NS-Hha/YdgT binds at a nucleation site (left); H-NS-Hha has a higher DNA binding affinity than H-NS-YdgT (see figure 3 -figure supplement 1d). Next, individual H-NS dimers or H-NS-Hha/YdgT assemblies multimerize and give rise to long protein-DNA filaments (right). Whereas YdgT association enhances the cooperativity of H-NS-DNA filament formation, Hha largely abolishes the cooperativity in H-NS-DNA filament formation.

